## Supplementary material for "*Drosophila* model to clarify the pathological significance of OPA1 in autosomal dominant optic atrophy": Table 1

| Figure no. | genotype | notes |
|---|---|---|
| 1B | GMR-Gal4/UAS-dOPA1-HA; UAS-Mito-mCherry/+ | |
| 1C, E | GMR-Gal4/40D-UAS; UAS-MitoGFP/+ | control |
| 1D, E | GMR-Gal4/+; UAS-MitoGFP/UAS-dOPA1 RNAi (BDSC 32358) | dOPA1 RNAi |
| 1F, J | GMR-Gal4/40D-UAS | control |
| 1G, J | GMR-Gal4/+; UAS-dOPA1 RNAi (BDSC 32358)/+ | dOPA1 RNAi |
| 2A, C | GMR-Gal4/40D-UAS; UAS-MitoGFP/+ | control |
| 2B, C | GMR-Gal4/+; UAS-MitoGFP/UAS-dOPA1 RNAi (BDSC 32358) | dOPA1 RNAi |
| 2D, F, H | GMR-Gal4/40D-UAS; UAS-mitoQC (BDSC 91641)/+ | control |
| 2E, G, H | GMR-Gal4/+;UAS-mitoQC (BDSC 91641)/UAS-dOPA1 RNAi (BDSC 32358) | dOPA1 RNAi |
| 3B, D, G, I, F, K | GMR-Gal4/40D-UAS | control |
| 3C, E, H, J, F, K | GMR-Gal4/+; UAS-dOPA1 RNAi (BDSC 32358)/+ | dOPA1 RNAi |
| 4A, D | FRT42D/ FRT42D, w+, cl/; ey-Gal4, UAS-flp/+ | control |
| 4B, D | FRT42D, dOPA1[s3475]/ FRT42D, w+, cl/; ey-Gal4, UAS-flp/+ | dOPA1 clone |
| 4C, D | UAS-dOPA1, FRT42D, opa1[s3475]/ FRT42D, w+, cl/; ey-Gal4, UAS-flp/+ | rescue |
| 5B-D | Tub-Gal80TS/+; Tub-Gal4/UAS-HA-hOPA1-myc | hOPA1 wt |
| 5B-D | Tub-Gal80TS/+; Tub-Gal4/UAS-HA-hOPA1[I382M]-myc | I382M |
| 5B-D | Tub-Gal80TS/+; Tub-Gal4/UAS-HA-hOPA1[D438V]-myc | D438V |
| 5B-D | Tub-Gal80TS/+; Tub-Gal4/UAS-HA-hOPA1[R445H]-myc | R445H |
| 5B-D | Tub-Gal80TS/+; Tub-Gal4/UAS-HA-hOPA1[2708del]-myc | 2708del |
| 5E | FRT42D/ FRT42D, w+, cl/; ey-Gal4, UAS-flp/+ | control |

| | | |
|---|---|---|
| 5E | FRT42D, dOPA1[s3475]/ FRT42D, w+, cl/; ey-Gal4, UAS-flp/+ | dOPA1 clone |
| 5E | UAS-dOPA1-HA, FRT42D, opa1[s3475]/ FRT42D, w+, cl/; ey-Gal4, UAS-flp/+ | dOPA1 rescue |
| 5E | FRT42D, dOPA1[s3475]/ FRT42D, w+, cl/; ey-Gal4, UAS-flp/UAS-HA-hOPA1-myc | hOPA1 rescue |
| 5E | FRT42D, dOPA1[s3475]/ FRT42D, w+, cl/; ey-Gal4, UAS-flp/UAS-HA-hOPA1[I382M]-myc | I382M rescue |
| 5E | FRT42D, dOPA1[s3475]/ FRT42D, w+, cl/; ey-Gal4, UAS-flp/UAS-HA-hOPA1[D438V]-myc | D438V rescue |
| 5E | FRT42D, dOPA1[s3475]/ FRT42D, w+, cl/; ey-Gal4, UAS-flp/UAS-HA-hOPA1[R445H]-myc | R445H rescue |
| 5E | FRT42D, dOPA1[s3475]/ FRT42D, w+, cl/; ey-Gal4, UAS-flp/UAS-HA-hOPA1[2708del]-myc | 2708del rescue |
| 5F | UAS-dicer2/+; GMR-Gal4/40D-UAS; UAS-Mito-GFP/UAS-luciferase RNAi (BDSC 31603) | control |
| 5F | UAS-dicer2/+; UAS-dOPA1-RNAi (VDRC 106290)/GMR-Gal4; UAS-Mito-GFP/UAS-luciferase RNAi (BDSC 31603) | dOPA1 knockdown |
| 5F | UAS-dicer2/+; UAS-dOPA1-RNAi (VDRC 106290)/GMR-Gal4; UAS-Mito-GFP/UAS-HA-hOPA1-myc | hOPA1 rescue |
| 5F | UAS-dicer2/+; UAS-dOPA1-RNAi (VDRC 106290)/GMR-Gal4; UAS-Mito-GFP/UAS-HA-hOPA1[I382M]-myc | I382M rescue |
| 5F | UAS-dicer2/+; UAS-dOPA1-RNAi (VDRC 106290)/GMR-Gal4; UAS-Mito-GFP/UAS-HA-hOPA1[D438V]-myc | D438V rescue |
| 5F | UAS-dicer2/+; UAS-dOPA1-RNAi (VDRC 106290)/GMR-Gal4; UAS-Mito-GFP/UAS-HA-hOPA1[R445H]-myc | R445H rescue |

| | | |
|---|---|---|
| 5F | UAS-dicer2/+; UAS-dOPA1-RNAi (VDRC 106290)/GMR-Gal4; UAS-Mito-GFP/UAS-hOPA1[2708del]-myc | 2708del rescue |
| 6A | GMR-Gal4/40D-UAS | contrrol |
| 6A | GMR-Gal4/+; UAS-HA-hOPA1-myc/+ | hOPA1 wt |
| 6A | GMR-Gal4/+; UAS-HA-hOPA1[I382M]-myc/+ | I382M |
| 6A | GMR-Gal4/+; UAS-HA-hOPA1[D438V]-myc/+ | D438V |
| 6A | GMR-Gal4/+; UAS-HA-hOPA1[R445H]-myc/+ | R445H |
| 6A | GMR-Gal4/+; UAS-HA-hOPA1[2708del]-myc/+ | 2708del |
| 6B | ey3.5flp/+; FRT42D, dOPA1[s3475]/FRT42D, W+, cl; UAS-HA-hOPA1-myc/GMR-GAl4, UAS-HA-hOPA1-myc | hOPA1 wt + hOPA1 wt in dOPA1 clone |
| 6B | ey3.5flp/+; FRT42D, dOPA1[s3475]/FRT42D, W+, cl; UAS-HA-hOPA1-myc/GMR-GAl4, UAS-HA-hOPA1[I382M]-myc | hOPA1 wt + I382M in dOPA1 clone |
| 6B | ey3.5flp/+; FRT42D, dOPA1[s3475]/FRT42D, W+, cl; UAS-HA-hOPA1-myc/GMR-GAl4, UAS-HA-hOPA1[D438V]-myc | hOPA1 wt + D438V in dOPA1 clone |
| 6B | ey3.5flp/+; FRT42D, dOPA1[s3475]/FRT42D, W+, cl; UAS-HA-hOPA1-myc/GMR-GAl4, UAS-HA-hOPA1[R445H]-myc | hOPA1 wt + R445H in dOPA1 clone |
| 6B | ey3.5flp/+; FRT42D, dOPA1[s3475]/FRT42D, W+, cl; UAS-HA-hOPA1-myc/GMR-GAl4, UAS-HA-hOPA1[2708del]-myc | hOPA1 wt + 2708del in dOPA1 clone |
| S1B | GMR-Gal4/40D-UAS; UAS-MitoGFP/+ | control |
| S1C | GMR-Gal4/+; UAS-mitoGFP/UAS-dOPA1 RNAi (BDSC 32358) | dOPA1 RNAi |
| S1D | GMR-Gal4/+; UAS-mitoGFP/UAS-Milton RNAi (BDSC 44477) | Milton RNAi |
| S2A-B | GMR-Gal4/40D-UAS | control |
| S2A-B | GMR-Gal4/UAS-dOPA1 RNAi (VDRC 106290) | dOPA1 RNAi |
| S3A-B | GMR-Gal4/40D-UAS | control |
| S3A-B | GMR-Gal4/+; UAS-dOPA1 RNAi (BDSC 32358)/+ | dOPA1 RNAi#1 |
| S3A-B | GMR-Gal4/UAS-dOPA1 RNAi (VDRC 106290) | dOPA1 RNAi#2 |
| S4A-D | UAS-dicer2/+; Tub-Gal80TS/40D-UAS ; Tub-Gal4/UAS-luciferase RNAi (BDSC 31603) | control |
| S4A-D | UAS-dicer2/+; Tub-Gal80TS/UAS-dOPA1-RNAi (VDRC 106290) ; Tub-Gal4/UAS-luciferase RNAi (BDSC 31603) | dOPA1 knockdown |
| S4A-D | UAS-dicer2/+; Tub-Gal80TS/UAS-dOPA1-RNAi (VDRC 106290) ; Tub-Gal4/UAS-HA-hOPA1-myc | hOPA1 rescue |

| | | |
|---|---|---|
| S4A-D | UAS-dicer2/+; Tub-Gal80TS/UAS-dOPA1-RNAi (VDRC 106290) ; Tub-Gal4/UAS-HA-hOPA1[I382M]-myc | I382M rescue |
| S4A-D | UAS-dicer2/+; Tub-Gal80TS/UAS-dOPA1-RNAi (VDRC 106290) ; Tub-Gal4/UAS-HA-hOPA1[D438V]-myc | D438V rescue |
| S4A-D | UAS-dicer2/+; Tub-Gal80TS/UAS-dOPA1-RNAi (VDRC 106290) ; Tub-Gal4/UAS-HA-hOPA1[R445H]-myc | R445H rescue |
| S4A-D | UAS-dicer2/+; Tub-Gal80TS/UAS-dOPA1-RNAi (VDRC 106290) ; Tub-Gal4/UAS-HA-hOPA1[2708del]-myc | 2708del rescue |
| S6A | UAS-dicer2/+; UAS-dOPA1-RNAi (VDRC 106290)/GMR-Gal4; UAS-Mito-GFP, UAS-HA-hOPA1-myc/ UAS-luciferase RNAi (BDSC 31603) | dOPA1 knockdown |
| S6A | UAS-dicer2/+; UAS-dOPA1-RNAi (VDRC 106290)/GMR-Gal4; UAS-Mito-GFP, UAS-HA-hOPA1-myc/ UAS-HA-hOPA1-myc | hOPA1 wt + hOPA1 wt in dOPA1 knockdown |
| S6A | UAS-dicer2/+; UAS-dOPA1-RNAi (VDRC 106290)/GMR-Gal4; UAS-Mito-GFP, UAS-HA-hOPA1-myc/UAS-HA-hOPA1[I382M]-myc | hOPA1 wt + I382M in dOPA1 knockdown |
| S6A | UAS-dicer2/+; UAS-dOPA1-RNAi (VDRC 106290)/GMR-Gal4; UAS-Mito-GFP, UAS-HA-hOPA1-myc/UAS-HA-hOPA1[D438V]-myc | hOPA1 wt + D438V in dOPA1 knockdown |
| S6A | UAS-dicer2/+; UAS-dOPA1-RNAi (VDRC 106290)/GMR-Gal4; UAS-Mito-GFP, UAS-HA-hOPA1-myc/UAS-HA-hOPA1[R445H]-myc | hOPA1 wt + R445H in dOPA1 knockdown |
| S6A | UAS-dicer2/+; UAS-dOPA1-RNAi (VDRC 106290)/GMR-Gal4; UAS-Mito-GFP, UAS-HA-hOPA1-myc/ UAS-hOPA1[2708del]-myc | hOPA1 wt + 2708del in dOPA1 knockdown |
| S6B-E | UAS-dicer2/+; Tub-Gal80TS/UAS-dOPA1-RNAi (VDRC 106290) ; Tub-Gal4, UAS-HA-hOPA1-myc/ UAS-luciferase RNAi (BDSC 31603) | dOPA1 knockdown |
| S6B-E | UAS-dicer2/+; Tub-Gal80TS/UAS-dOPA1-RNAi (VDRC 106290) ; Tub-Gal4, UAS-HA-hOPA1-myc/ UAS-HA-hOPA1-myc | hOPA1 wt + hOPA1 wt in dOPA1 knockdown |
| S6B-E | UAS-dicer2/+; Tub-Gal80TS/UAS-dOPA1-RNAi (VDRC 106290) ; Tub-Gal4, UAS-HA-hOPA1-myc/ UAS-HA-hOPA1[I382M]-myc | hOPA1 wt + I382M in dOPA1 knockdown |
| S6B-E | UAS-dicer2/+; Tub-Gal80TS/UAS-dOPA1-RNAi (VDRC 106290) ; Tub-Gal4, UAS-HA-hOPA1-myc/ UAS-HA-hOPA1[D438V]-myc | hOPA1 wt + D438V in dOPA1 knockdown |
| S6B-E | UAS-dicer2/+; Tub-Gal80TS/UAS-dOPA1-RNAi (VDRC 106290) ; Tub-Gal4, UAS-HA-hOPA1-myc/ UAS-HA-hOPA1[R445H]-myc | hOPA1 wt + R445H in dOPA1 knockdown |
| S6B-E | UAS-dicer2/+; Tub-Gal80TS/UAS-dOPA1-RNAi (VDRC 106290) ; Tub-Gal4, UAS-HA-hOPA1-myc/ UAS-HA-hOPA1[2708del]-myc | hOPA1 wt + 2708del in dOPA1 knockdown |
| S7A-B | GMR-Gal4/40D-UAS | control |
| S7A-B | GMR-Gal4/+; UAS-dOPA1 K273A/+ | dOPA1 dominant negative form |
